# Quasi-Equilibrium State Based Quantification of Biological Macromolecules in Single-Molecule Localization Microscopy

**DOI:** 10.1101/2023.07.17.549270

**Authors:** Xuecheng Chen, Yaqian Li, Xiaowei Li, Jielin Sun, Daniel M. Czajkowsky, Zhifeng Shao

## Abstract

The stoichiometry of molecular components within supramolecular biological complexes is often an important property to understand their biological functioning, particularly within their native environment. While there are well established methods to determine stoichiometry *in vitro*, it is presently challenging to precisely quantify this property *in vivo*, especially with single molecule resolution that is needed for the characterization stoichiometry heterogeneity. Previous work has shown that optical microscopy can provide some information to this end, but it can be challenging to obtain highly precise measurements at higher densities of fluorophores. Here we provide a simple approach using already established procedures in single-molecule localization microscopy (SMLM) to enable precise quantification of stoichiometry within individual complexes regardless of the density of fluorophores. We show that by focusing on the number of fluorophore detections accumulated during the quasi equilibrium-state of this process, this method yields a 50-fold improvement in precision over values obtained from images with higher densities of active fluorophores. Further, we show that our method yields more correct estimates of stoichiometry with nuclear pore complexes and is easily adaptable to quantify the DNA content with nanodomains of chromatin within individual chromosomes inside cells. Thus, we envision that this straightforward method may become a common approach by which SMLM can be routinely employed for the accurate quantification of subunit stoichiometry within individual complexes within cells.

## Introduction

Determining the stoichiometry of biomolecular components within macromolecular assemblies is critical for an understanding of how they function. Many different biophysical methodologies have been employed to this end, including size-exclusion chromatography, analytical ultracentrifugation, gel electrophoresis, mass spectrometry, and optical spectroscopy [1]. However, while highly effective, these methods cannot be readily used for *in situ* studies, where such complexes interact and function with other biomolecules within their native environment [2]. In this regard, optical microscopy, and in particular super-resolution microscopic methods such as single-molecule localization microscopy (SMLM), can provide *in situ* structural information at resolutions down to the size of individual macromolecular complexes (∼20 nm) [3]. With these methods, images are produced by acquiring thousands of conventional, diffraction-limited images that each contain a very sparse distribution of fluorescent molecules, followed by fitting of their positions at a high precision. For densely labeled targets, such sparsity can be obtained with photoswitchable fluorophores or transiently binding labels [4]. With some complexes that have well-separated subunits (or sub-complexes), such as the nuclear pore complexes (NPCs), a resolution of ∼20 nm alone is sufficient to identify distinct subunits within individual complete complexes, providing at least some information about the underlying molecular stoichiometry [5]. However, most large macromolecular complexes are not structured in this way and details of the stoichiometry are thus not readily obtainable at the presently attainable resolutions. In these situations, the only alternative to obtain information about the stoichiometry of complexes is by analysis of the number of detections of the fluorescent molecules, and indeed previous SMLM studies determined stoichiometry based on the total number of detections measured in each locus [6]. However, the accuracy and precision for such counting is significantly dependent on the density of the fluorophores in each diffraction-limited region. In particular, for highly dense labeling (which is often advantageous for higher-resolution images), it can be challenging to accurately determine the number of fluorophores at the initial stages of imaging, as there is a preponderance of fluorophores in the on-state, which can lead to a mischaracterization of multiple detections as owing to a single fluorophore and thus an imprecise measure of the calculated stoichiometry [7]. With photo-activation localization microscopy (PALM), it is possible to artificially introduce a much lower density of fluorescent proteins through transfection, and then infer stoichiometry by analyzing the number of observed localizations through feature extraction of the blinking events and then fitting to a kinetic model [8, 9], or via Markov modeling [10], Bayesian inference [11], or neural network modeling [12]. These approaches have indeed been successfully applied to examine low-oligomeric protein organization in super-resolved protein clusters [13− 15]. However, by requiring such transfections to produce controlled low protein densities, PALM poses technical challenges that may not be suitable for many proteins, and moreover, could not be used for studies of nucleic acid quantification. Further, such a strategy could not readily be adopted with immunolabeling procedures, which are commonly used by many other SMLM methods, as this may result in non-uniform or loss of labeling [16].

Here we provide a simple alternative for quantifying proteins and DNA with stochastic optical reconstruction microscopy (STORM), a commonly used SMLM method. Our method avoids issues with mis-allocation of detections at high densities of fluorophores, especially at the beginning of image acquisition, by instead focusing on the imaging period during which only a low density of fluorophores is active within any one frame [17]. This imaging period reflects a “quasi-equilibrium” state of the (photo-converting) fluorophores, and appears to be a common characteristic of fluorescent molecules, such as Alexa 647 fluorophores [17]. We show that our method exhibits a 50-fold lower variation in detection counts for both microtubules and NPCs than otherwise found at higher densities of fluorophores, and is, moreover, adaptable to quantify the extent of EdU-labeled DNA within individual chromatin domains in human fibroblast cell nuclei. As our method requires no different imaging strategy than is presently employed by most STORM investigations, we anticipate that it will be readily adopted by other groups as a straightforward procedure by which the stoichiometry of biomolecular assemblies can be easily determined *in situ*.

## Materials and methods

### 1. Cell lines, antibodies and chemicals

The COS-7 (African green monkey kidney fibroblast-like cell line) and U2OS cells were purchased from the National Collection of Authenticated Cell Cultures (China), whereas the BJ cells (Human fibroblast cell line) were purchased from ATCC.

The monoclonal anti-alpha tubulin antibody was purchased from Sigma-Aldrich (T6199), while the monoclonal anti-Nup133 antibody was purchased from Abcam (ab181355). The Alexa 647-conjugated secondary antibodies were purchased from ThermoFisher (A21235 and A21244).

F-ara-EdU, thymidine and hydroxyurea were purchased from Sigma-Aldrich. The DMEM medium and other chemicals used for cell culturing were purchased from ThermoFisher (Gibco brand). Fetal bovine serum was purchased from BI (04-001-1A).

### 2. Cell culture

For immunostaining of microtubules and nuclear pore complexes, the COS-7 cells and U2OS cells were cultured on 10-cm dishes (Corning, 430591) at 37 °C and 5% CO2 in DMEM medium with L-glutamine (ThermoFisher, 11965092) containing 10% FBS (BI, 04-001-1A), penicillin (100 U/ml) and streptomycin (0.1 mg/ml) (ThermoFisher, 15140122). Exponential growing cells were seeded onto glass-bottom dishes (Cellvis, D29-10-1.5N) at a ∼30% density and grown for 12 h prior to fixation.

For chromatin DNA imaging, the BJ cells were cultured in the same way as described above. For synchronization, cells were seeded onto 10-cm dishes with a ∼30% density and cultured for 18 h with pre-warmed fresh medium containing 2 mM thymidine (Sigma-Aldrich, 89270). The cells were next washed three times with pre-warmed 1xPBS and then incubated in pre-warmed fresh medium with no thymidine for 8 h. Cells were trapped at the G1/S boundary by incubating the cells in media containing 1 mM hydroxyurea (Sigma-Aldrich, H8627) for 18 h. Pre-warmed medium containing 10 μM F-ara-EdU (Sigma-Aldrich, T511293) was then added to the synchronized cells. The F-ara-EdU medium was discarded after 12 h as the DNA replication had finished and fresh medium without EdU was added to the cells, which were allowed to grow for 5 to 7 generations to dilute the EdU to single chromosomes. The cells were then seeded onto the glass-bottom dishes at a ∼ 30% density and again synchronized to the G1/S boundary as described above before fixation. The calibration sample was prepared as the same in our previous work [18]. In short, the EdU-incorporated DNA was purified from the cells treated with the same procedures as above, followed by fixation to the poly-L-lysine (Sigma-Aldrich, P8920) treated cover-glasses for 30 min.

### 3. Immunostaining and click reaction

The microtubules were labeled in the COS-7 cells. In short, the cells were washed three times with 1xPBS and fixed for 10 minutes with 4% paraformaldehyde (Sigma-Aldrich, 158127) and 0.1% glutaraldehyde (Sigma-Aldrich, 340855) in buffer-A (buffer-A, 10 mM MES, 150 mM NaCl, 5 mM EGTA, 5 mM glucose and 5 mM MgCl2, pH 6.1), followed by quenching with freshly prepared 0.1% NaBH4 (Sigma-Aldrich, 452882) for another 7 min. We next permeabilized and blocked the cells for non-specific binding by incubating with 1xPBS containing 0.5% Triton X-100 and 5% BSA for 1 h. The microtubules were labeled using mouse anti-alpha tubulin antibody (1:1000 diluted in 1xPBS containing 5% BSA) for 1 hour followed by three washes (5 min each) with 0.1% Tween-20 in 1XPBS. Next the cells were labeled with the Alexa 647-conjugated secondary antibody (1: 500) for another 1 h followed by washing with 0.1% Tween-20 in 1XPBS.

The nuclear pore complexes were labeled in U2OS cells. The cells were incubated for 40 s with 0.2% Triton X-100 and then immediately fixed with 2.4% paraformaldehyde in 1xPBS for 10 minutes, followed by 10 min incubation with 0.1 M glycine. The permeabilization and non-specific blocking were achieved following the same as the procedure for microtubules. The rabbit anti-Nup133 antibody (1:100 in 1% BSA) was then incubated with the cells for 1.5 hour followed by extensive washing and labeling with Alexa 647-conjugated secondary antibody (1: 500) for 1 hour.

For the labeling of chromatin DNA, cells with F-ara-EdU incorporated DNA were washed and fixed with 4% paraformaldehyde for 10 min, followed by the same quenching, permeabilization and blocking process as above. The Cu(I)-catalyzed azide-alkyne cycloaddition (CuAAC) reaction was then performed to add Azido-Alexa 647 fluorophores (ThermoFisher, C10340) to the F-ara-EdU loci on the chromatin DNA. In short, the cells were incubated for 2 h with 10 nm Azido-Alexa 647 fluorophores in clicking buffer (100 mM Tris-HCl, 100mM sodium L-ascorbate and 2 mM CuSO4, pH 7.5), followed by washing the cells three times with 1x PBS containing 1 mM EDTA. The calibration samples were clicked following the same process.

### 4. SMLM imaging (Nikon N-STORM)

SMLM imaging was performed using a Nikon N-STORM microscope, equipped with a 200 mW 647 nm laser; an APO 100x, NA 1.49 oil objective and an Andor iXon3 EMCCD camera with a pixel size of 16 μm. The imaging buffer consisted of 0.5 mg/ml glucose oxidase (Sigma-Aldrich, G2133), 40 μg/ml catalase (Sigma-Aldrich, C40), 10% (w/v) glucose (Sigma-Aldrich, G8270) and 143 mM βME (Sigma-Aldrich, M6250) dissolved in TN buffer (50 mM Tris-HCl, pH 8.0 and 10 mM NaCl).

100, 000 SMLM raw Images were taken with a 20 ms exposure time, 30x EM gain and 5x conversion gain for microtubules and 255x EM gain and 1x conversion gain for nuclear pore complexes, respectively. Super-resolution images were reconstructed using the software ThunderSTORM [19].

### 5. SMLM Data processing

We processed the data following a similar procedure as described previously [17] using ThunderSTORM [19]. In short, we set a threshold of 4 times the standard deviation of the fluctuations of the detected background signal for the detection of the molecules. Molecules that appear consecutively in more than one frame at the same position (< 20 nm shift) were merged into one detection (or fluorescence switching event). The number of detections associated with any molecule was calculated by summing the detections within a window of 100 s (that is, 5000 frames). Based on our measurements as well as prior work [17], we identified defined the quasi-equilibrium state switching regime as the imaging period between 400 to 600 s for the Alexa 647 fluorophore used in this work. The total number of detections was determined by summing the measured detections from all molecules that did not photobleach during the experiment.

## Results

### 1. Basic principle of our quantification method

In SMLM experiments, fluorophores switch reversibly between a fluorescent on- and nonfluorescent off-state. Defining the rates of the transitions from the on-to the off-state and from the off-to the on-state as k_off_ and k_on_, respectively, it can be shown that the number of single molecules in the on-state N_F(t)_ changes with time t according to [7]:

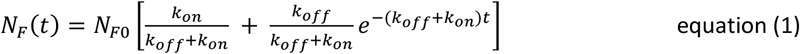

Here, N_F0_, the number of fluorophores in the on-state at t=0, is also the total number of fluorescent molecules, since in typical experiments, all of the photo-switchable fluorophores are initially in their on-state [7]. It can be seen that after a sufficiently long period of time (t → ∞),

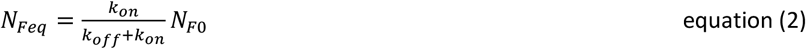

That is, the number of fluorophores in the on-state in this quasi-equilibrium state is a fixed fraction of the total number of fluorescent molecules, dependent on the inherent properties of that fluorophore (k_off_ and k_on_).

Confirming this expectation, Zhuang and colleagues reported that most of the fluorophores that are typically used in STORM were found to achieve such a quasi-equilibrium state between 400 to 600 s during image acquisition [17]. Further, they showed that a very high fraction (more than 80%) of the fluorophores remain active at this time (that is, they are not photobleached) and that the density of on-state molecules during this period is generally sparse enough such that the overwhelming majority of events in a single diffraction-limited area are single molecules events [17]. For example, with a frame rate of 1/20 ms^-1^, there are ∼ 5 switching events in total for one Alexa-647 fluorophore during this 200 s/10000 frame window [17]. This is the key property that we take advantage of to quantify stoichiometry with our method: In most cases, the density of fluorophores at initial illumination is quite high (as required for maximal labeling of molecular components) such that more than one fluorophore is fluorescent within each diffraction-limited area. This is a well-known concern in SMLM studies, which results in false-positive localizations and imaging artifacts [7]. With regards to the determination of stoichiometry, this issue fundamentally prevents an accurate measure of N_F0_. We also note that, in actual experiments, it is also impossible to ensure all the molecules are, in fact, fluorescent at initial illumination. Thus, by focusing on a temporal regime in which it is most likely that only one molecule per diffraction-limited area is in the on-state at any moment, we can obtain a more faithful measure of the number of molecules.

Under our imaging conditions, with a frame rate of 1/20 ms^-1^, each fluorophore is expected to be in the on-state for no more than a single frame. Thus, during analysis of the acquired data, molecules that appear consecutively in more than one frame at the same position (that is, within 20 nm) are conventionally merged together and counted as a single detection. During initial illumination when there are multiple fluorophores in the on-state within the same diffraction-limited area, these multiple fluorophores are merged together and not actually discerned as multiple detections (that is, one for each fluorophore). By contrast, during the quasi-equilibrium period, when there is most likely only one fluorophore per diffraction-limited area that is in the on-state in one frame, there is greater confidence that a measured detection is indeed owing to a single fluorophore. This is also the case when there are multiple fluorophores within the same resolution-limited localization – their number (that is, the stoichiometry) is most accurately determined when only one of the fluorophores is in the on-state in any one frame.

Thus, our method of determining the stoichiometry entails measuring the total number of detections associated with each resolution-limited localization during the quasi-equilibrium period between 400 to 600 s, and then dividing this by the number of detections within this same time period for a single fluorophore. The latter can be obtained, for example, from a separate experiment of a low surface density of the fluorophores. Such a measurement is necessary since the number of detections within this period (determined by k_on_) is generally different for different fluorophores.

### 2. Quantification of tubulin density with microtubules based on analysis of the quasi-equilibrium regime

To validate this approach and show that it leads to more precise measurements of stoichiometry, we examined the changes in detection with time with two different biological macromolecules, microtubules and NPCs. For the former, we labeled COS-7 cells with a monoclonal anti-alpha-tubulin antibody and an Alexa 647-conjugated secondary antibody (Figure 1A). As shown in Figure 1B, the temporal behavior of the detection density, measured from 15 microtubule filaments, is well-described according to theoretical expectations (equation (1)). From this, we found that the average number of accumulated detections in total is 2357 ± 182 per micron (n=15), while that in the quasi-equilibrium interval is 384 ± 0.67 per micron (n=15) (Figure 1B). The relative standard deviation of the latter (0.67/384) is thus ∼50 times smaller than the former (182/2357). Hence, there is a substantial improvement in the precision of the detection quantification by focusing on the quasi-equilibrium interval.

**Figure 1.**
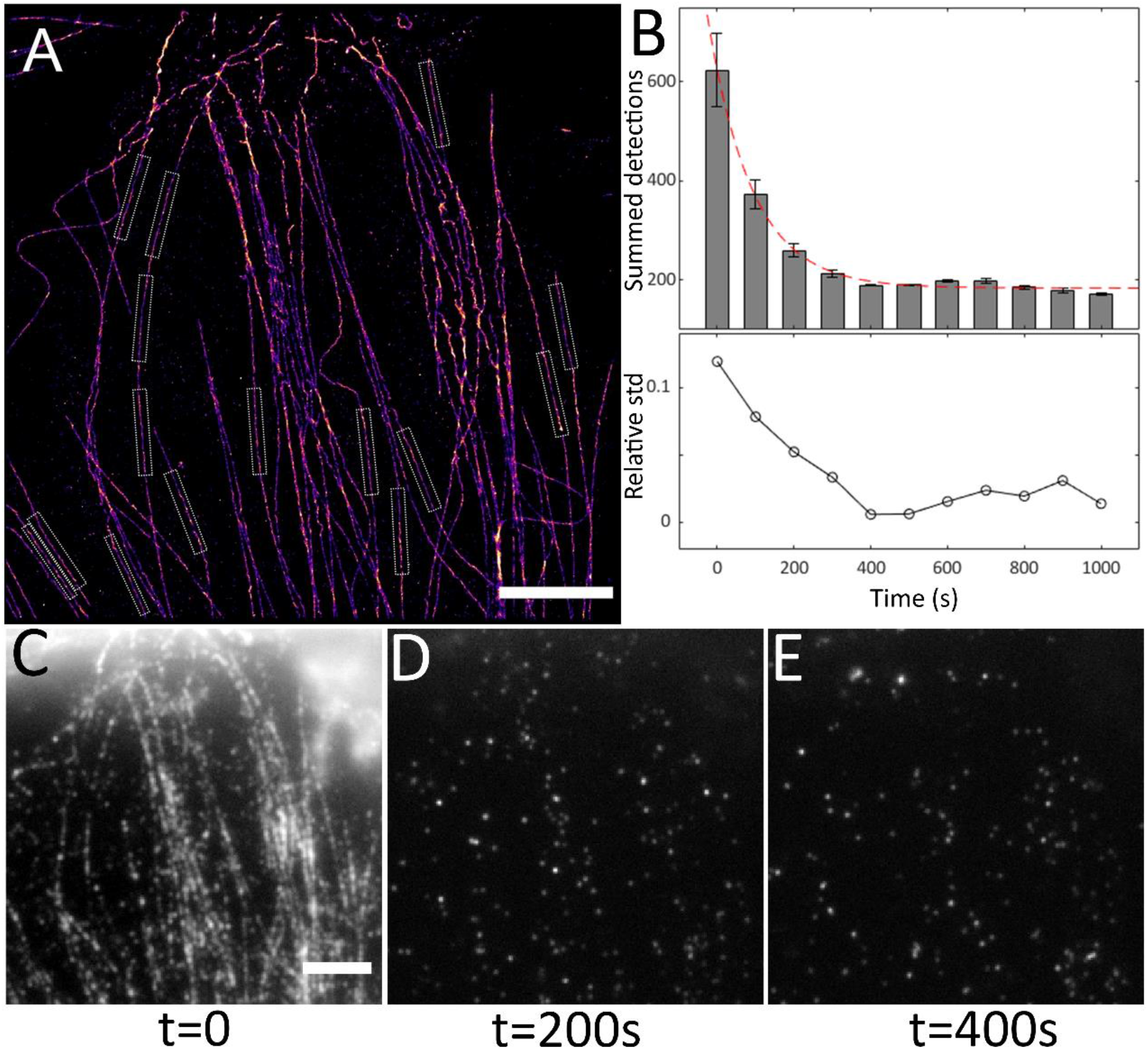
SMLM data of microtubules in COS-7 cells showing the changes in the number of detections with time. (A) Reconstructed SMLM image of the microtubules. The boxed regions depict examples of the segments of the microtubules within which the number of detections were counted. (B) The upper panel depicts the time evolution of the detection counts per micron along the microtubules (n=15), and the lower panel is the relative standard deviation of these measurements. It can be seen that, during the time period between 400 to 600 s, the system attains a quasi-equilibrium state in which the relative standard deviation in detection count is minimal. (C-E) Select frames from the SMLM data showing the high density of on-state fluorophores at earlier time points. Scale bar, A, C_E: 5 μm.

Previous experiments of single Alexa 647 fluorophores measured ∼5 detections during this quasi-equilibrium time period [17]. Since, according to the supplier and our previous results [18], there are 3 Alexa 647 fluorophores per antibody on average, we thus expected that each Alexa 647-bound secondary antibody would yield ∼15 detections within this quasi-equilibrium time interval. This number is in excellent agreement with the detection measurements obtained from individual foci (spots) beside the microtubules (Supplementary figure S1A), which exhibited a value of 17.1 ± 2.0 (Supplementary figure S1C). Based on this number, we calculate that the average labeling density of the microtubules is (384/15) ∼ 26 antibodies per micron. Given the typical size of a primary and secondary antibody (∼ 15 nm) and that of a tubulin dimer (∼ 8 nm), we expected that a closely-packed distribution of antibody labeling would yield a secondary antibody for every 3 to 4 tubulin dimers along the tubule axis. This value is close to our measured value of one every ∼5 dimers, in good agreement with the nearly continuous distribution of labeling apparent in these images.

### 3. Determination of the stoichiometry of nuclear pore complexes using quasi-equilibrium state-based quantification

As a further validation, we next examined NPCs in U2OS cell, labeling with a monoclonal anti-Nup133 antibody and an Alexa 647-conjugated secondary antibody (Figure 2A). In many images, the resolution of the images was indeed sufficient to identify individual subunits, which thereby enabled a more direct validation of the calculated stoichiometry with our approach. We measured both the total number of detections and the number of detections within the quasi-equilibrium period of 100 Nup133 rings. As with the microtubules described above, the temporal decay of the detection number is well-described by theoretical expectations (equation (1)). Further, we found that the relative standard deviation of this quasi-equilibrium detection count (0.28/158) is also ∼50 times smaller than that for the total detection count (172/2057) (Figure 2B), thus confirming the increased precision in this measurement. We then measured the number of detections for 300 well-resolved individual subunits of the NPCs, and found a similar 50-fold smaller variation in the precision of the quasi-equilibrium count (Supplementary figure S2). Based on these measurements, using values obtained with the total number of detections, one would conclude (2057/184) ∼ 11 subunits for the complete NPC ring, whereas with values obtained from the quasi-equilibrium period, we find (158/21) ∼ 7.5 subunits, in good agreement with the known octameric stoichiometry of this complex [5].

**Figure 2.**
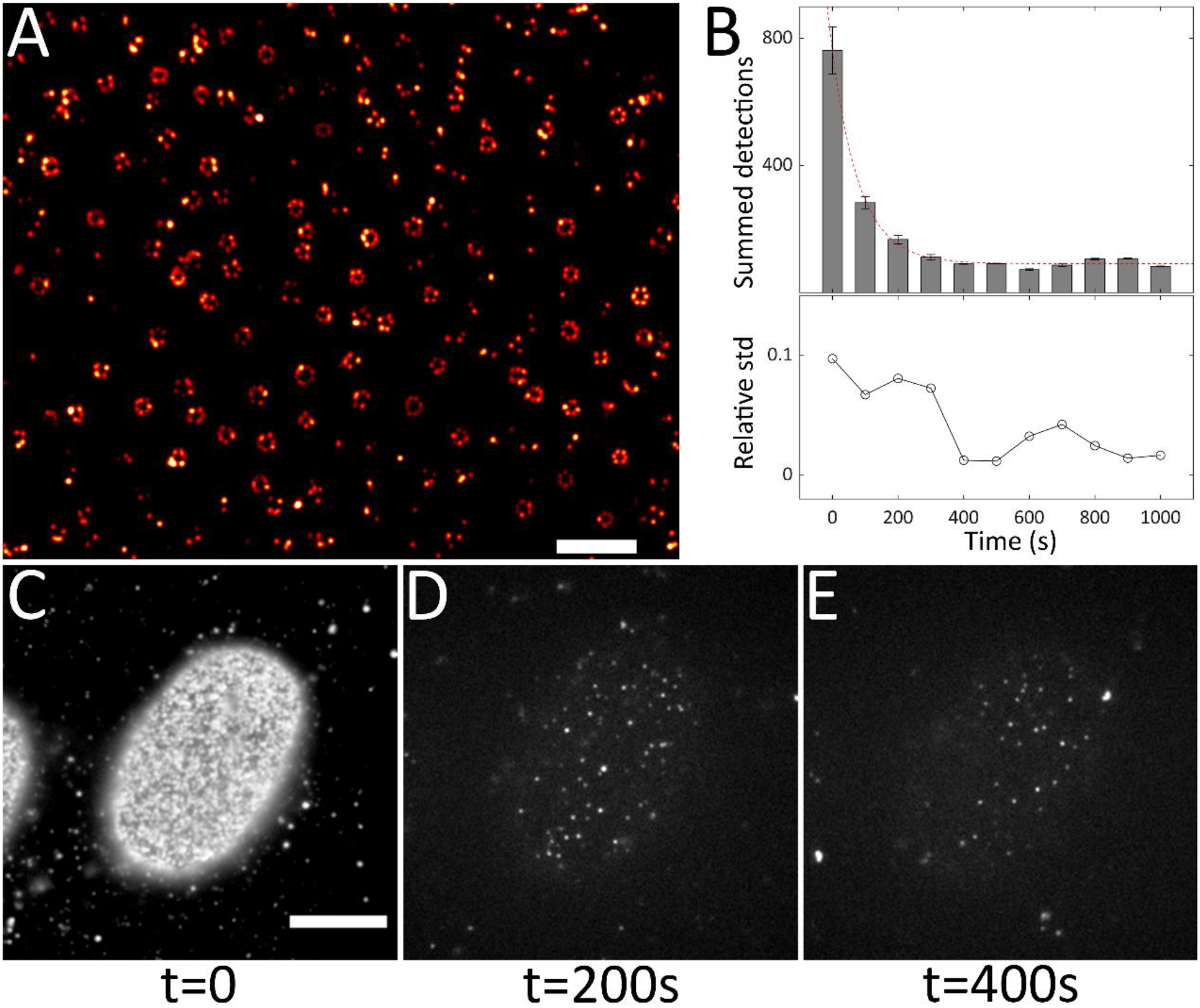
SMLM data of nuclear pore complexes (Nup133) in U2OS cells. (A) Typical SMLM image of the nuclear pore complexes. (B) The upper panel is the temporal evolution of the number of detections within complete Nup133 rings (n=100), and the lower panel is the relative standard deviation of these measurements. (C-E) Individual frames of the SMLM data showing the differences in the number of on-state fluorophores at different time points. Scale bar, A: 0.5 μm; C-E: 5 μm.

### 4. Quantification of DNA content within in vivo chromatin nanodomains

Finally, we sought to apply our method for the quantification of the DNA density within individual chromatin domains in cells. Following our previous method to study chromatin structures with STORM using EdU and click chemistry [18], we firstly blocked fibroblast cells at G1/S phase by double-thymidine synchronization, and then released them into S phase in the presence of 10 μM EdU for one generation (resulting the incorporation of 1 EdU per ∼300 bp [18]). To determine the number of detections per length of genomic DNA, we purified 1 kb (∼3 EdU) and 2 kb (∼6 EdU) EdU-DNA fragments from sonicated genomic DNA. After adsorption of the DNA on a coverslip, we performed click chemistry with Alexa 647-conjugated azide followed by SMLM imaging (Figure 3A). The temporal change in detection number followed that expected from equation (1) (Figure 3B), from which we determined a number within the quasi-equilibrium period to be about 7.1 ± 0.5 per kb (Figure 2C). Since it is known that the typical click chemistry reaction results in a bleaching of ∼50% of the fluorophores [21], this measure of detection number is in good agreement with the expected 15 detections for 3 EdUs (clicked with Alexa-647 fluorophores) per 1 kb DNA based on the previous results for single Alexa-647 fluorophores (5) [9]. With this calibration, we then repeated the aforementioned labeling scheme but then washed away the EdU after the first cell cycle and allowed the cells to continue to grow for another 5 to 7 generations. In this way, chromosomes labeled during the first generation become diluted into different cells, ultimately resulting in cells with only a single EdU-labeled chromosome that we earlier showed resulted in a significantly improved resolution with STORM [18]. Indeed, imaging such samples revealed cells with one or two labeled regions per cell, each exhibiting heterogeneous nanodomains, identified as described previously [18] (Figure 2D-F). With our calibration, we found that the imaged loci in Figure 2G is about 100 Mb, consisting of ∼ 500 nanodomains that consist of between 2 to 10 kb DNA (Figure 2H-J), which is consistent with previous work [22-24].

**Figure 3.**
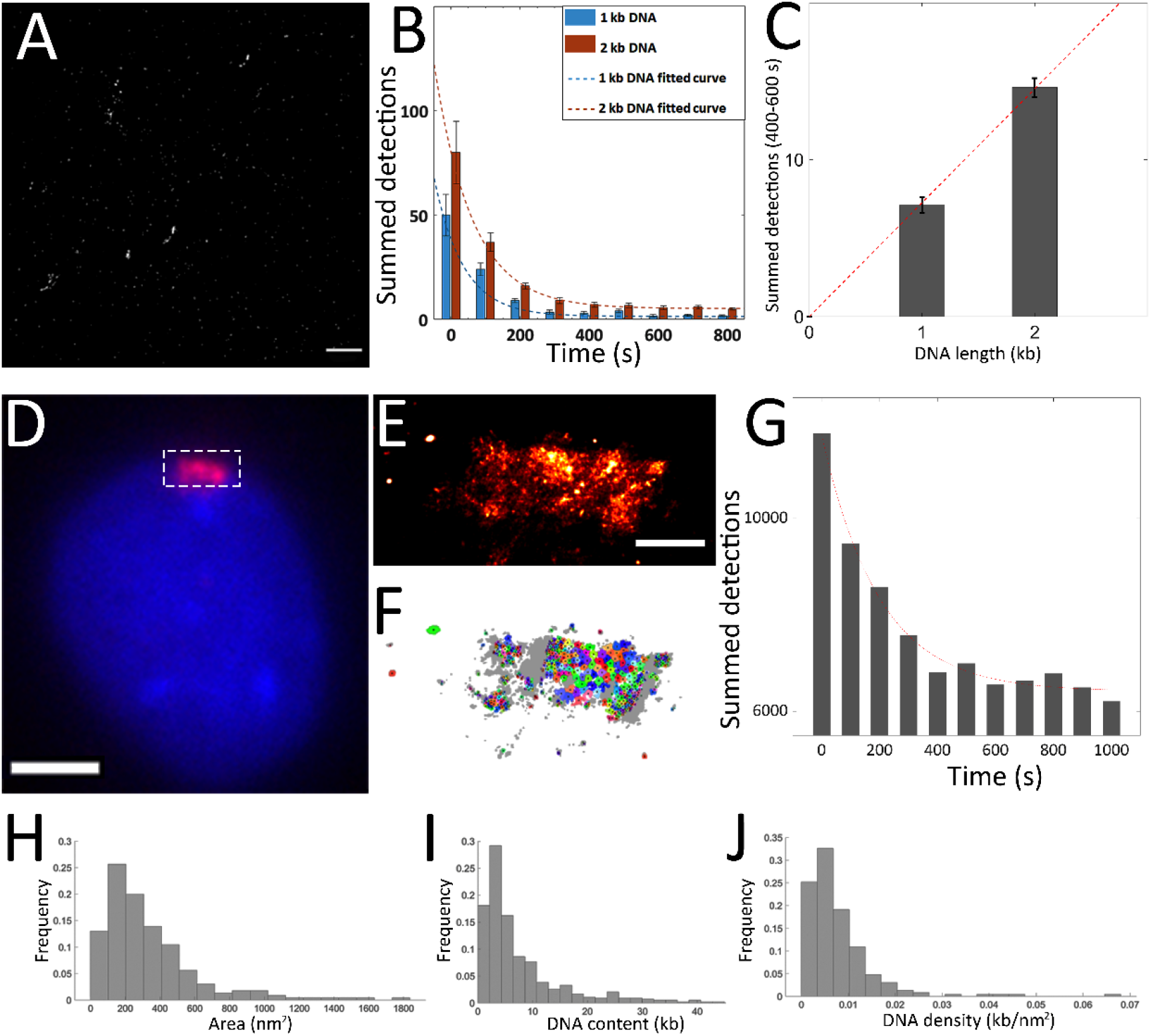
Quantification of chromatin DNA based on SMLM data within the quasi-equilibrium period. (A) SMLM image of 1-kb genomic EdU-DNA clicked with Alexa-647 fluorophores on cover glass. (B) Temporal evolution of detection number from individual foci of 1-kb and 2-kb genomic EdU-DNA. (C) The number of detections per kb for the two samples within the quasi-equilibrium period. (D) Wide-field view of a single labeled chromosome with all DNA labeled with DAPI (blue) and the single EdU-incorporated chromosome labeled with Alexa-647 (red). (E) SMLM image of the single chromosome in (D). (F) Clustering results from the SMLM image identifying nanodomains of chromatin. (G) Time course of the detection number from all loci within a single chromosome. (H-J) Distribution of area, DNA density and DNA content of individual nanodomains, with the latter two measurements based on data during the quasi-equilibrium period. Scale bar, A: 2 μm; D: 5 μm and E: 1 μm.

## Discussion

In conclusion, we have developed a simple method to precisely quantify the stoichiometry of proteins and DNA in SMLM studies based on the number of detections in the quasi-equilibrium interval during imaging. By comparing the differences in the number of detections of different complexes, it should be possible to obtain a measure of the fold-differences in the relative stoichiometry between complexes without any calibration or prior-knowledge of the targets. However, with appropriate calibration, as shown here, our method can also be used for the determination of the absolute stoichiometry of the molecular components within individual complexes. Such information is ultimately essential to understand the structures and, as importantly, the heterogeneity in the structures of such assemblies under native conditions that underlies their functioning within the cell.

## Acknowledgments

We would like to thank Mr. Ming Cheng for his assistance in preparation of this manuscript. The authors acknowledge support from Zhiyuan Innovative Research Center (ZIRC) of Shanghai Jiao Tong University. The authors are also grateful for the generous support from Nikon Instruments Co, Ltd. (Tokyo, Japan)

