## Supplementary Materials for "Quasi-Equilibrium State Based Quantification of Biological Macromolecules in Single-Molecule Localization Microscopy"

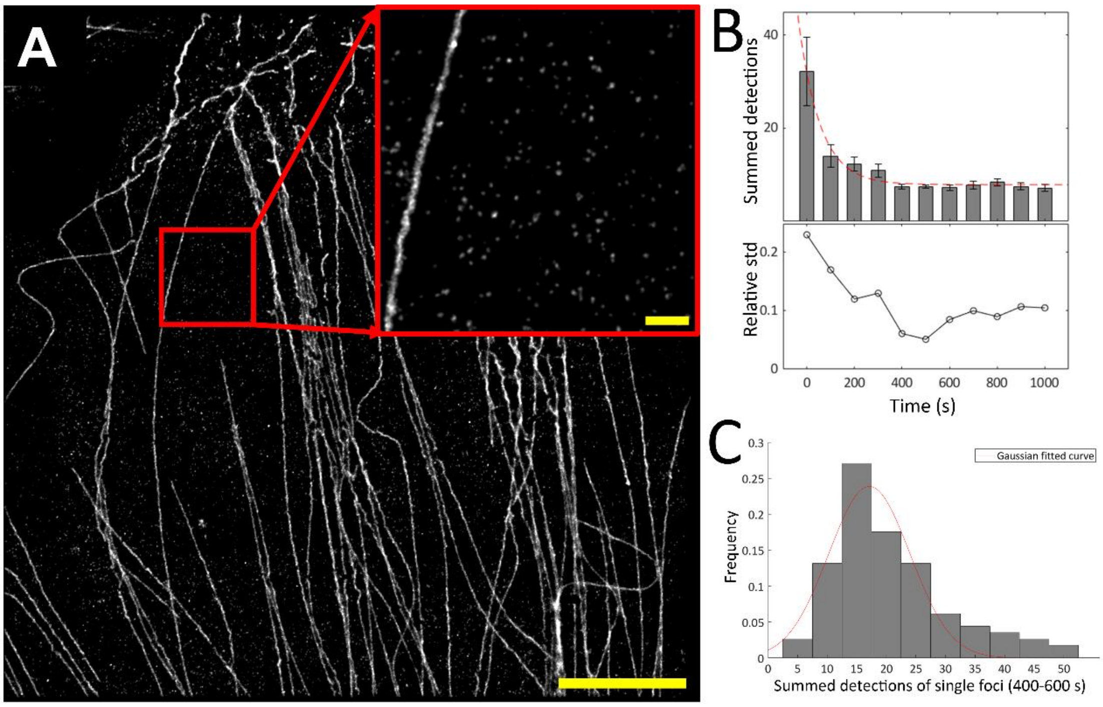


**Supplementary Figure S1.** Analysis of single foci in the SMLM reconstructed images of microtubules. (A) Reconstructed SMLM images with an inset showing numerous single foci with a similar intensity, which we suggest are individual secondary antibodies. (B) The average number of detections from the single foci in (A) (n=50). (C) Histogram of the quasi-equilibrium state detections of the single foci (n=200) in (A) and the Gaussian fitted curve showing the mean quasi-equilibrium state detection count of 17.1$\pm$2.0. Scale bar, A: 5 µm; A-inset: 0.5 µm;

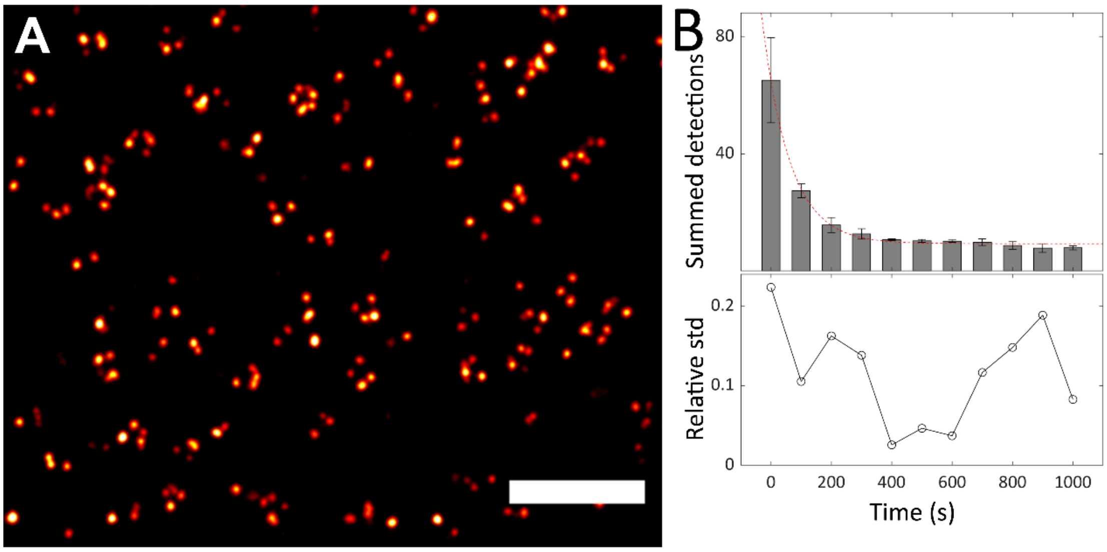


**Supplementary Figure S2.** Analysis of single subunits of nuclear pore complexes. (A) SMLM image showing single Nup133 subunits. (B) Temporal evolution of the summed detections from isolated Nup133 subunits (n=200). Scale bar, A: 0.5 µm
